# Subthalamic nucleus controls cautious action timing under threat

**DOI:** 10.64898/2025.12.30.696140

**Authors:** Ji Zhou, Muhammad S. Sajid, Sebastian Hormigo, Manuel A. Castro-Alamancos

## Abstract

Adaptive goal-directed actions performed under the threat of punishment introduce a demand for caution, often expressed as delayed response timing that balances urgency against error risk by allowing more time for cognitive evaluation. Although this form of temporal regulation is essential for survival, its underlying neural mechanisms remain poorly understood. We show that glutamatergic neurons in the subthalamic nucleus (STN) regulate the timing of cued actions to avoid harm. Optogenetic activation of the STN, or its projections to the midbrain but not the globus pallidus, modulates action timing in a frequency-dependent manner, accelerating initiation to the extent that animals can no longer respond cautiously, defer actions, or stop initiated actions. These results show that the STN shapes goal-directed behavior by gating action-initiation timing, establishing its midbrain projections as a key circuit for balancing urgency and caution under threat.

**Significance statement:** This work shows that activity in the subthalamic nucleus (STN) helps set the timing of actions triggered by warning cues. By activating STN neurons and their projections to the midbrain, we found that stimulation made animals respond faster, and high-frequency activation eliminated their ability to delay actions when caution was required or to withhold actions that would otherwise be punished. These findings highlight a key brain pathway that governs when actions are initiated, a function that is often disrupted in neurological and psychiatric disorders.

## Introduction

Adaptive actions in threatening environments hinge on deciding when to act, balancing urgency against the risk of making mistakes. When a cue signals that failing to act will be punished, but the same action leads to punishment under other conditions (e.g., when the cue is absent or when a different cue is presented), animals must respond quickly enough to avoid harm while ensuring that acting is appropriate. Cautious responding by slowing action initiation provides extra time for sensory processing and cognitive evaluation, reducing the likelihood of costly errors. Delayed response timing has long been recognized as a behavioral signature of additional cognitive processing, since Donders’s work in the mid-19^th^ century (Donders, 1969; Sternberg, 1969; Posner, 2005). Although speed–accuracy tradeoffs are well characterized in reward-based decision-making, far less is known about how the brain controls action timing when aversive outcomes are possible (Smith and Ratcliff, 2004; van Maanen et al., 2011; Guitart-Masip et al., 2012; Ratcliff and Frank, 2012; Yee et al., 2022).

The subthalamic nucleus (STN) is a glutamatergic nucleus in the basal ganglia that integrates cortical and pallidal inputs and influences movement through its projections to the midbrain. STN activity is well known to correlate with movement (Isoda and Hikosaka, 2008; Callahan et al., 2024; Zhou et al., 2025), and has been linked to response slowing (Frank et al., 2007; Cavanagh et al., 2014; Herz et al., 2024) as well as action cancellation in humans (Aron, 2011). In rodents, some studies indicate that STN initiates or facilitates movement (Watson et al., 2021; Fan et al., 2023; Friedman and Yin, 2023), whereas others indicate movement suppression (Fife et al., 2017; Guillaumin et al., 2021; Xie et al., 2022). Recently, we showed that STN plays a critical role in generating cued actions under threat, encoding the cautious responses required to avoid harm and enabling their performance (Zhou et al., 2025). Here, we investigated whether STN controls the timing of cued actions under threat by optogenetically modulating STN activity and its principal ascending and descending projections.

## Results

### STN excitation drives movements in the ipsiversive direction

In a previous study, we found that nearly half of recorded STN neurons were significantly modulated by movement with subsets encoding movement onset and direction (Zhou et al., 2025). Approximately one quarter of neurons showed strong activation during contraversive turns, whereas another quarter exhibited weaker ipsiversive biases. Here we tested whether optogenetic excitation of STN neurons would causally bias movement toward contraversive or ipsiversive directions. We expressed ChR2 in the STN of Vglut2-Cre mice using bilateral injections of a Cre-inducible AAV and implanted a dual optical fiber bilaterally (STN-ChR2 mice, n = 11 mice; Fig. 1A). During open-field exploration (Fig. 1B), we applied blue light trains (continuous or 1-ms pulses at 10, 20, 40, and 66 Hz for 2 seconds) to excite STN neurons unilaterally or bilaterally at varying levels. Light trains were delivered randomly at different intervals within the same sessions. These procedures were also performed in No Opsin control mice (n = 6), which did not express ChR2.

**Figure 1.**
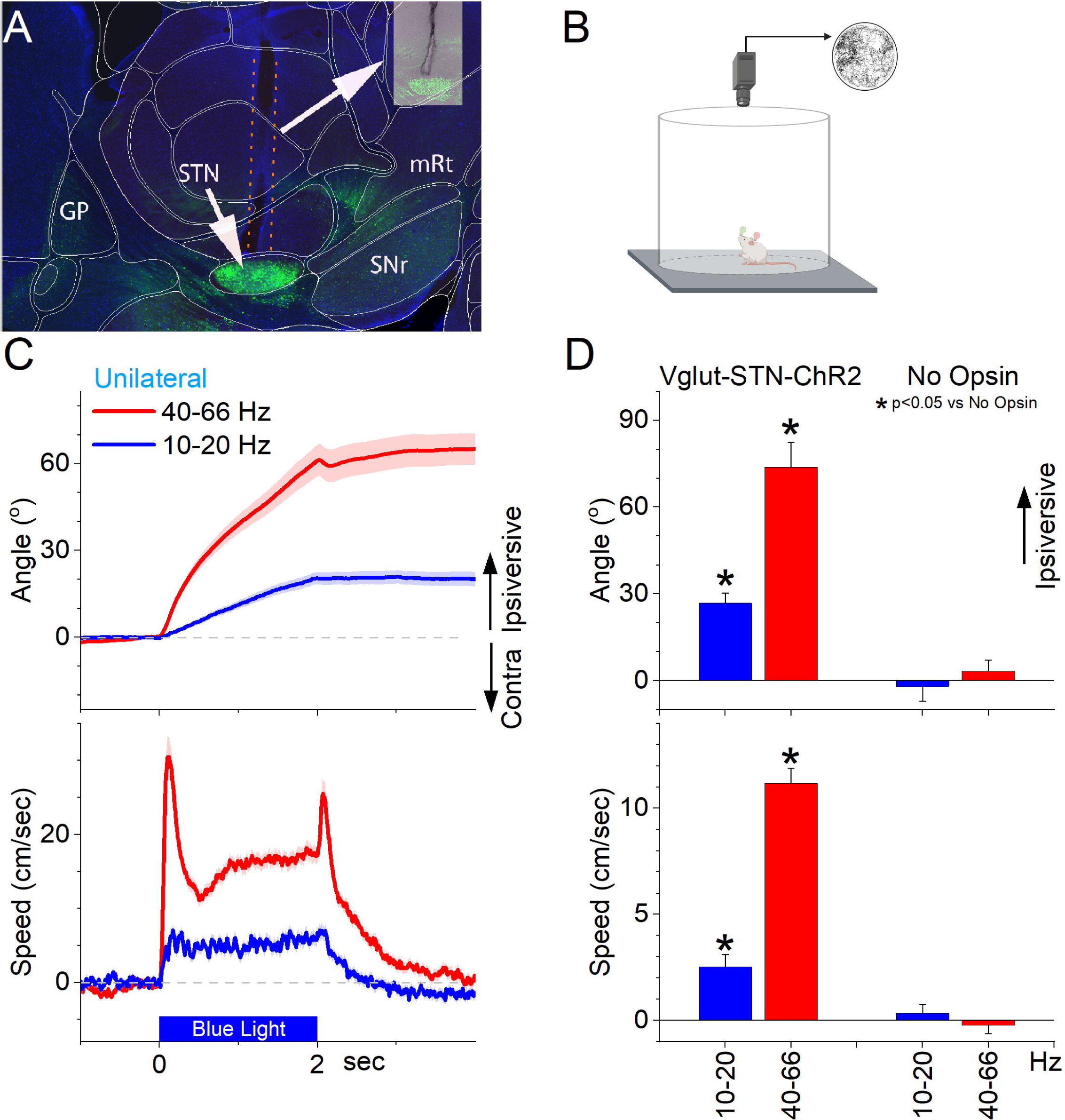
Optogenetic activation of ChR2-expressing STN glutamatergic neurons produces an ipsiversive turning bias. ***A***, Parasagittal section showing the optical fiber tract reaching STN and ChR2 fluorescence expressed in glutamatergic neurons. The section was aligned with the Allen brain atlas. |***B***, Schematic showing the experiment arena. ***C***, Unilateral optogenetic STN excitation drives an ipsiversive head orienting bias. Light patterns (2 sec at different frequencies) are delivered randomly as mice explore an arena. The top panel shows head orienting angle change from the onset of the light. The bottom panel shows the movement speed. Note the strong ipsiversive bias during 40-66 Hz STN excitation. ***D***, Population data comparing Opsin mice expressing ChR2 in STN neurons with No Opsin mice. Asterisks denote significant differences (p<0.05) between Opsin and No Opsin mice.

Unilateral optogenetic excitation of STN neurons produced a robust ipsiversive head orienting direction bias, with the strength of the effect depending on the frequency of blue light stimulation (Fig. 1C). Even the lowest effective stimulation frequencies and powers elicited ipsiversive movements. There was a significant effect of light frequency on directional bias in the STN-ChR2 mice (Fig. 1C,D; Mixed ANOVA Tukey t(60)=8.6 p<0.0001, 10-20 Hz vs 40-66 Hz) but not in the NoOpsin mice (Tukey t(60)=0.7 p=0.94). There was also a significant difference in directional bias between STN-ChR2 and No Opsin mice for 10-20 Hz (Fig. 1C,D; Tukey t(60)=4.6 p=0.009) and 40-66 Hz (Tukey t(60)=11.3 p<0.0001) blue light. Light applied as continuous 2 s pulses (not shown) was also tested and produced similar but stronger effects than the high frequency trains.

The effects of bilateral optogenetic excitation are further examined below; however, when compared directly within the STN-ChR2 group in the open field arena, bilateral stimulation induced significantly more movement than unilateral stimulation at both 10–20 Hz (2Way ANOVA Tukey t(37)=3.3 p=0.02 Unilateral vs Bilateral) and 40-66 Hz (Tukey t(37)=8.4 p<0.0001). The higher frequencies caused more movement than the lower frequencies when applied either unilaterally (Tukey t(37)=7.3 p<0.0001) or bilaterally (Tukey t(37)=12.3 p<0.0001).

Together, these findings indicate that STN excitation promotes movement in a frequency-dependent manner. Although subsets of STN neurons encode contraversive or ipsiversive directions during spontaneous behavior (Zhou et al., 2025), unilateral stimulation predominantly elicits ipsiversive movements (Fig. 1C,D). This dissociation suggests that while some STN neurons encode contraversive movements, they may not directly drive them, pointing to a more nuanced role for the STN in modulating movement direction.

### STN excitation controls cued actions

In our previous study, we showed that STN neurons are critically involved in generating and robustly encode cued actions under threat, particularly the cautious responses required to avoid punishment (Zhou et al., 2025). The critical role of the STN in generating cued actions under threat stands in contrast to related regions, including other basal ganglia nuclei (Hormigo et al., 2021b, 2021a; Zhou et al., 2024), the prefrontal cortex (Zhou et al., 2022), and the neighboring zona incerta (Hormigo et al., 2020, 2023), which modulate these behaviors but are not required for producing them. Here, we used optogenetic excitation to test whether modulating STN glutamatergic activity causally alters these behaviors as mice performed the same sequence of cued procedures in a shuttle box (AA1–AA3; Fig. 2A,B). In AA1, mice avoid an aversive US by shuttling between compartments (action) when CS1 was presented during a 7 s avoidance interval, performing a high percentage of correct actions (active avoids) and few errors (escapes triggered by the US starting at 7 s from CS onset). CS1 contingencies were identical across AA1-3. In AA2, the only change versus AA1 is that spontaneous intertrial crossings (ITCs) during the intertrial interval (25-45 s) were punished by a short US, reliably shifting CS1 avoid latencies to longer values, consistent with increased caution (Zhou et al., 2022). AA3 introduced a challenging discrimination, where mice continued to actively avoid in response to CS1 but were required to withhold the action during CS2 to passively avoid a brief US (ITCs were no longer punished); crossings during CS2 in AA3 are signaled passive avoid errors, whereas ITCs in AA2 are unsignaled passive avoid errors.

**Figure 2.**
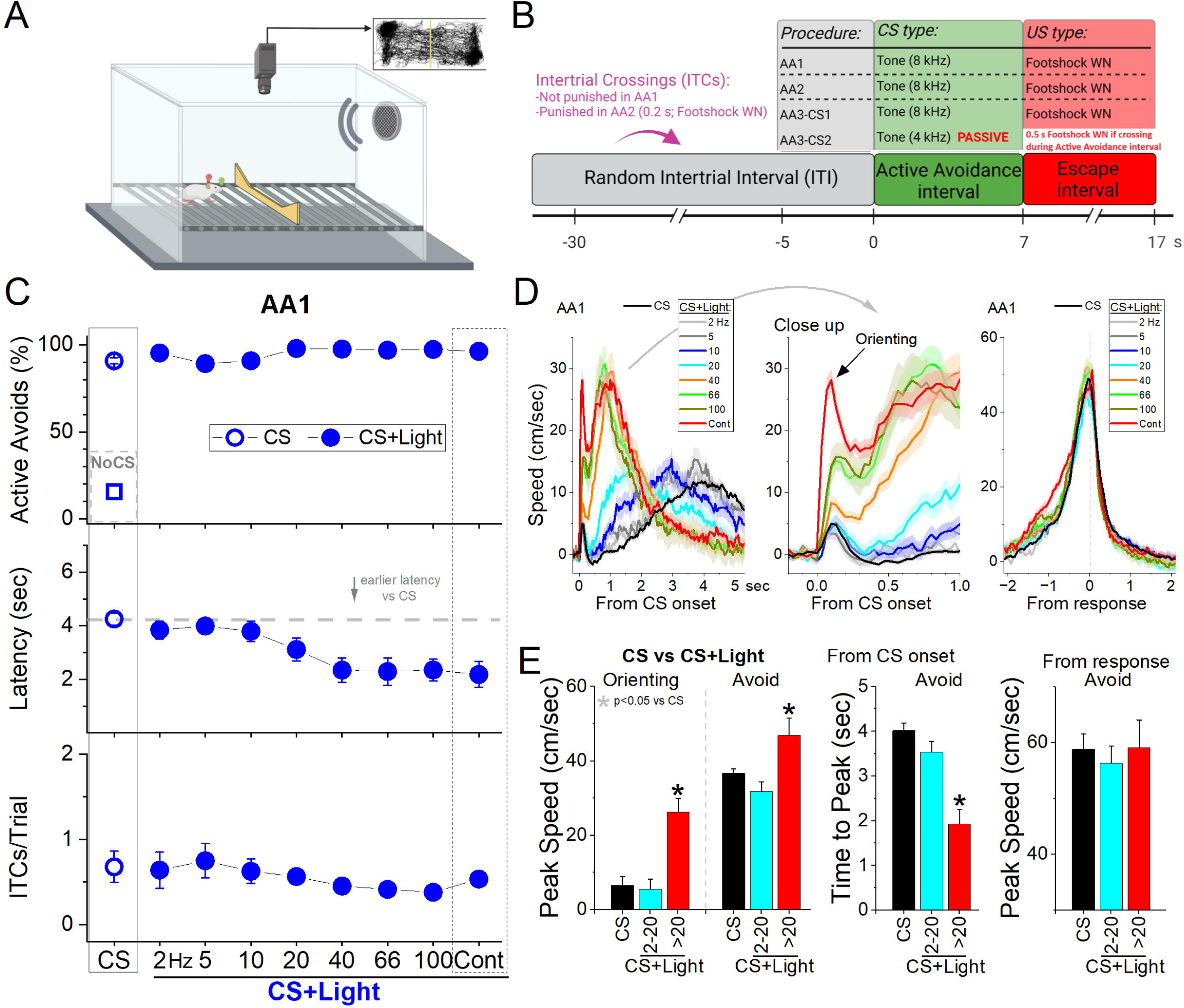
Optogenetic activation of ChR2-expressing STN glutamatergic neurons drives signaled active avoidance and can substitute for the natural CS. ***A***, Schematic showing the shuttle box used for signaled avoidance experiments. ***B,*** Schematic highlighting the trial types involved in AA1, AA2, and AA3 procedures. Trials consist of a cued (CS) avoidance interval when crossing (shuttling) avoids the escape interval when the US (footshock and white noise) is presented until mice escape (all mice escape <2.5 s). All trials are followed by a random intertrial interval when crossings are punished only in AA2, which is the only difference vs AA1. In AA3, CS2 trials are cued by a different CS that signals to not cross (passive avoidance). If mice cross during CS2 they are punished (0.5 s US). ***C,*** Effects of blue light patterns on signaled active avoidance. Blue circles show the effect of CS+Light trials (filled blue circles) compared to control CS trials (open blue circle) and catch NoCS trials (open blue squares). Note the shortening of the latency for higher frequencies, including Cont light. ***D***, Traces of the overall speed during AA1 for CS trials and CS+Light trials for different light patterns. The trials are aligned by CS onset (left and middle panels), which reveals the orienting response evoked by the CS followed by the avoidance action, or from avoid occurrence (right panel), which reveals the peak speed as the mice cross between compartments. ***E***, Population measures of peak speed and time to peak speed for CS and CS+Light trials from CS onset, for the orienting response and the avoidance response peaks from CS onset (left and middle panels). The right panel compares the peak speed measured from the avoid occurrence. CS+Light trials data are combined according to the light pattern into low frequency (2-20 Hz) or high frequency (>20 Hz and Cont). Asterisks denote significant differences (p<0.05) between CS trials and CS+Light trials.

To excite STN neurons at varying levels, we expressed ChR2 and applied a spectrum of blue light patterns in STN including trains (1-ms pulses at 2-100 Hz) and continuous (Cont) light. This approach has been noted for its value (González-Rueda and Tripodi, 2019) as it provides a graded modulation of neuronal activity, which we have characterized in several studies (e.g., (Hormigo et al., 2016, 2021a, 2021b)). Within each session, we compared randomly delivered CS (control), NoCS (catch), and CS+Light or Light (alone) trials. The light was delivered during the avoidance interval and continued during the escape interval if mice failed to avoid in active avoidance trials. Moreover, the light turns off when the animal avoids or escapes (shuttles). In addition, No Opsin mice were subjected to these same trials and compared to opsin expressing mice. Initially, we tested the effects of the optogenetic light (blue) in No Opsin mice (n=10) subjected to the behavioral procedures. In the absence of opsin expression, the light (CS vs CS+Light trials) had little effect on behavioral performance (active avoids rate and latency) during AA1 (RMAnova F(1,9)= 0.16 p=0.69; F(1,9)= 0.87 p=0.37), AA2 (RMAnova F(1,9)=2.65 p=0.14; F(1,9)= 0.08 p=0.78), or AA3 (2WayRMAnova F(1,9)= 0.01 p=0.9; F(1,9)= 0.00005 p=0.99) procedures.

We tested the effects of exciting glutamatergic STN neurons by expressing ChR2 in the STN of Vglut2-Cre mice with bilateral injections of a Cre-inducible AAV (Fig.1A, STN-ChR2, n=11 mice). Excitation of STN neurons during AA1 increased the percentage of active avoids (90.2±1.9 vs 97.2±1.5 %; CS vs CS+Light trials) but avoids are close to maximal in control CS trials and the effect only bordered significance (Fig. 2C circles top; Tukey t(5)= 3.6 p=0.051). However, STN excitation produced by blue light patterns >20 Hz, including Cont, sharply decreased avoid latencies (Fig. 2C circles middle; Tukey t(5)= 5.19 p=0.014). Movement tracking revealed that the effects of STN excitation (>20 Hz) on AA1 performance were associated with both a strong increase in the orienting response amplitude (Fig. 2D left; Tukey t(10)= 6.47 p=0.0027) and an earlier onset of the ensuing avoid, which peaked sooner measured from CS onset (Fig. 2D,E middle; Tukey t(10)= 9.94 p<0.0001). However, the peak speed of the avoids measured from the response occurrence was not affected by STN excitation (Fig. 2D,E right; Tukey t(10)= 0.36 p=0.96). This indicates that STN excitation leads to earlier onset avoids occurring at a similar peak speed as normal avoids. Therefore, instead of the characteristic avoid latencies that occur around the middle of the avoidance interval, avoids occur earlier as a function of STN excitation frequency without altering their peak speed.

To test if STN excitation alone could substitute for the natural CS, we employed Light trials. STN excitation at 10-20 Hz drove active avoids at percentage rates (Fig. 3A triangles top; Tukey t(10)= 2.55 p=0.21 CS vs Light trials 10-20 Hz) and latencies (Fig. 3A triangles middle; Tukey t(10)= 0.08 p=0.99) that did not differ from control CS trials indicating that excitation at these frequencies can generate normal avoidance responses. In contrast, STN excitation >20 Hz drove earlier onset avoid latencies compared to control CS trials (Fig. 3A triangles middle; Tukey t(10)= 4.76 p=0.017) or lower frequency (<20 Hz) STN excitation Light trials (Tukey t(10)= 4.84 p=0.016). Movement tracking revealed that STN excitation >20 Hz caused a strong increase in the orienting response (Fig. 3B,C; Tukey t(10)= 4.7 p=0.019), and an earlier avoid onset (Fig. 3B,C; Tukey t(10)= 5.42 p=0.0085), but did not change the peak speed of avoids compared to the CS (Fig. 3B,C; Tukey t(10)= 0.3 p=0.976). Thus, optogenetic STN activation driven by 1-ms blue light pulses at 10-20 Hz drives active avoids at normal speeds and onset latencies that are comparable to the active avoids driven by a natural CS, while stronger STN activation> 20 Hz shifts onset latencies earlier without altering peak speed.

**Figure 3.**
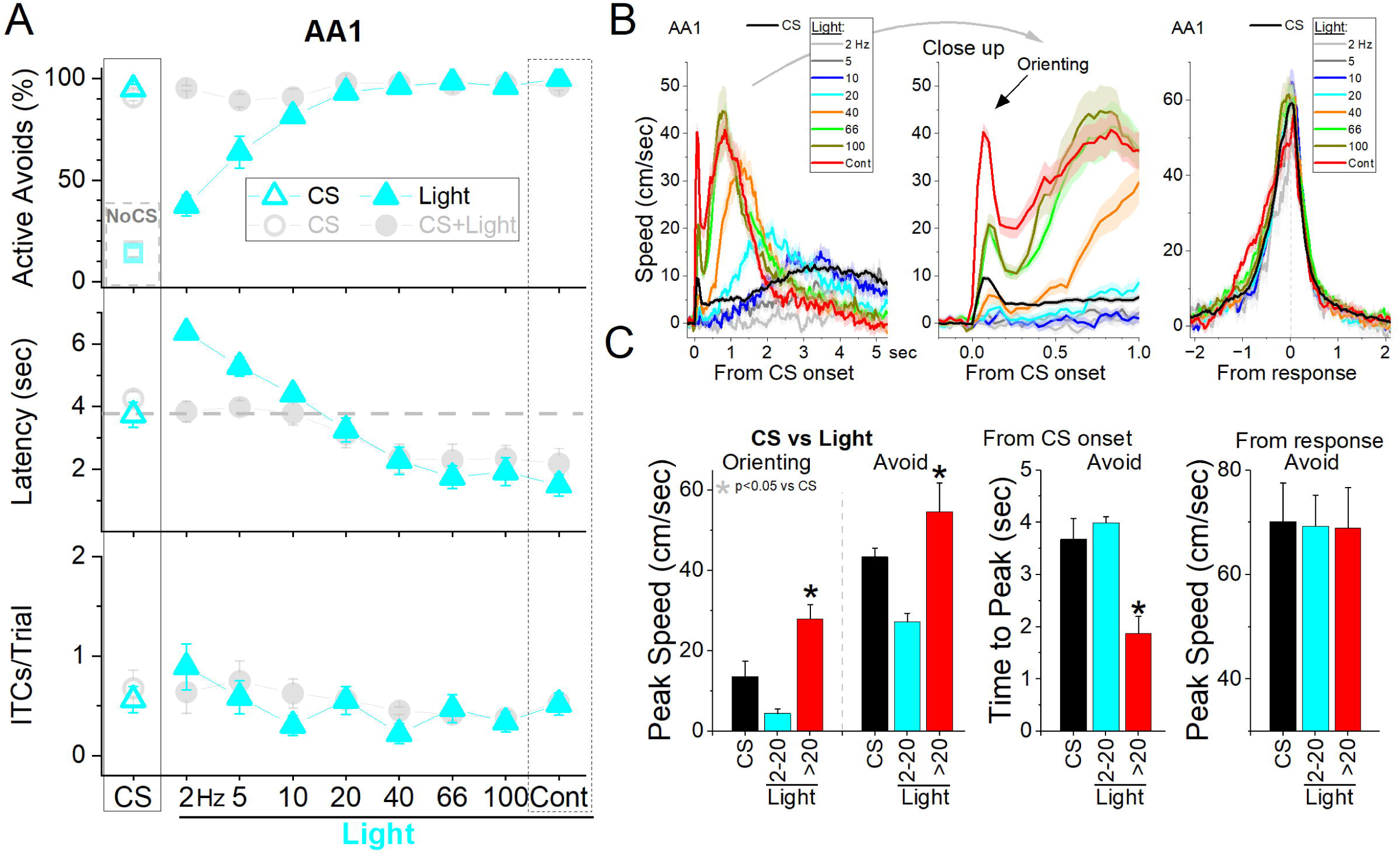
Optogenetic activation of ChR2-expressing STN glutamatergic neurons can substitute for the natural CS. ***A,*** Testing the ability of the light alone to serve as CS in the absence of the natural tone. Cyan triangles show the effect of Light trials (filled triangles) compared to control CS trials (open triangles) and catch NoCS trials (open squares). Note the shortening of the latency for higher frequencies including Cont light. The cyan triangles show the results for equivalent Light trials, where the CS is excluded, and the light serves as the CS. Light trials at low frequencies are ineffective at driving avoids. The deemphasized gray circles show the results of CS+Light trials from Figure 2 for comparison. ***B*,** Traces of the overall speed during AA1 for CS trials and Light trials for different light patterns. The trials are aligned by CS or Light onset (left and middle panels), which reveals the orienting response evoked by the CS followed by the avoidance action, or from avoid occurrence (right panel), which reveals the peak speed as the mice cross between compartments. ***C*,** Population measures of peak speed and time to peak speed for CS and CS+Light trials from CS onset, for the orienting response and the avoidance response peaks from CS onset (left and middle panels). The right panel compares the peak speed measured from the avoid occurrence. CS+Light trials data are combined according to the light pattern into low frequency (2-20 Hz) or high frequency (>20 Hz and Cont). Asterisks denote significant differences (p<0.05) between CS trials and CS+Light trials.

### STN excitation in naïve mice

The preceding Light trials occurred in trained mice. Therefore, we tested the effect of STN excitation in a group of naïve STN-ChR2 mice (n=5 mice) to determine if they would innately cross between compartments when STN excitation was applied even though it does not predict the US. During five sessions, we randomly presented NoCS (i.e., catch trials) and NoCS+Light trials (i.e., the light does not predict the US, which is not presented, but crossing stops the light just like an avoid stops the CS). Mice produced a high rate of crossings during STN excitation >20 Hz including Cont (Fig. 4A green circles; Tukey t(12)= 5.87 p=0.0063 NoCS vs NoCS+Light >20 Hz). To determine if during the naïve sessions the mice were learning to escape the STN stimulation over days, we compared the rate of crossings during the light between the first session and the fifth session but found no difference (Tukey t(4)= 0.6 p=0.6 Session 1 vs 5 for NoCS+Light >20 Hz). Thus, mice innately move away when STN is excited at high frequencies >20 Hz, which are the frequencies that drive early onset avoids when presented in CS+Light trials.

**Figure 4.**
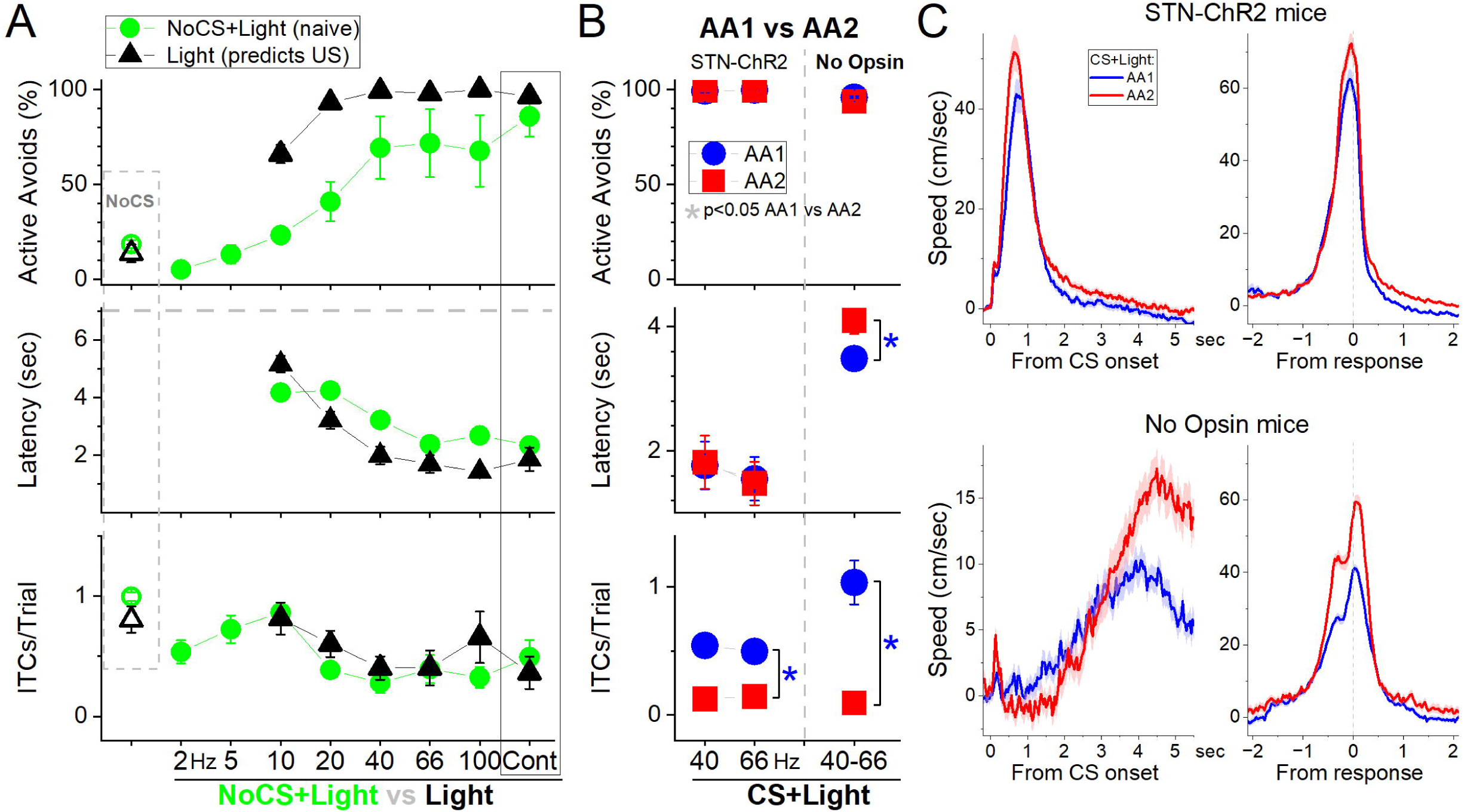
Optogenetic activation of ChR2-expressing STN glutamatergic neurons drives crossings in naïve mice and suppresses the development of caution. ***A,*** Effects of patterns of blue light applied in the STN of naïve mice where the light does not predict the US (NoCS+Light trials, green circles). This was followed by the addition of the US in regular Light trials (black triangles). Note that STN excitation in naïve mice drives a high rate of crossings. ***B***, Development of caution between AA1 and AA2 (by punishing ITCs) and reflected in the avoidance latency is suppressed by STN excitation in STN-ChR2 mice compared to No Opsin controls. The blue circles show AA1 sessions followed by red squares showing AA2 sessions in the same mice for STN-ChR2 and No Opsin mice. Note the abolishment of ITCs during AA2 and the shift of the latency longer only in No Opsin mice. Asterisks denote significant differences (p<0.05) between AA1 (blue) and AA2 (red). ***C,*** Traces of overall speed during AA1 and AA2 for the data shown in B. Note the characteristic changes in No Opsin mice are blunted by STN excitation in STN-ChR2 mice. This includes the rightward shift of the speed from CS onset, and the larger peak speed from response occurrence during AA2 compared to AA1.

On the sixth session, the NoCS+Light trials were converted to Light trials (i.e., the light now predicts the US; 2-5 Hz were excluded to reduce the number of punished trials). This led to an increase in avoidance rates for all STN excitation light patterns (Fig. 4A black triangles; 2WayRMAnova NoCS+Light vs Light trials F(1,16)=27.7 p=0.006) but this did not affect NoCS trials or ITCs (F(1,16)=0.58 p=0.48). In particular, Light trials at 10-20 Hz, which drove few crossings in naïve animals, became as effective as a natural CS at driving active avoids at normal latencies. Light trials >20 Hz were also effective at driving avoids but as for CS+Light trials they were characterized by early onset avoids. Furthermore, the addition of the natural CS to the Light trials (i.e., converting them to CS+Light trials; not shown) had no effect on avoidance rates for Light trials >5 Hz (Tukey t(32)<2.3 p>0.17 for 10-100 Hz and Cont), indicating that Light alone >5 Hz is equivalent to a natural CS, with 10-20 Hz matching the response features of normal avoids.

These results indicate that optogenetic STN activation at 10-20 Hz serves as a signal as effective as a natural CS in a cued goal-directed task while activation at higher frequencies drives earlier onset avoids without altering their peak speed and 2-5 Hz low frequency STN activation has little effect. STN activation has an intrinsic ability to serve as a CS during signaled active avoidance by associating with the US it predicts.

### STN excitation blocks the development of caution when errors are punished

During basic signaled active avoidance, which we term AA1, ITCs occur freely without consequence, but the succeeding AA2 procedure punishes ITCs, requiring mice to passively avoid during the intertrial interval. A characteristic, reliable feature of the transition between AA1 and AA2 procedures is a rightward shift of the cued active avoidance action to longer latencies in an apparent reflection of caution (Zhou et al., 2022). Since STN excitation shifts active avoid latencies earlier or leftward, we tested if STN excitation may interfere with the normal development of longer or rightward latency shifts between AA1 and AA2.

We used STN-ChR2 (n=6) and NoOpsin mice (n=7) subjected only to CS+Light trials (40-66 Hz; Fig. 4B) in successive AA1 sessions followed by AA2 sessions. During AA2, ITCs were rapidly abolished with little effect on active avoid rate, but avoidance latencies did not shift longer in STN-ChR2 mice, as they did in control No Opsin mice (Fig. 4B). Thus, the shift in avoid latency was larger in No Opsin mice compared STN-ChR2 mice subjected to STN excitation (F(1,11)=8.3 p=0.01; STN-ChR2 vs No Opsin). In fact, movement tracking revealed that STN excitation caused a leftward shift of speed during AA2 (Fig. 4C upper), which is earlier and opposite to what occurs in No Opsin (Fig. 4C lower) or normal mice (Zhou et al., 2022).

These results indicate that STN excitation >20 Hz is incompatible with the normal development of cautious behavior about generating a CS signaled action when the occurrence of the unsignaled action (ITC) is punished.

### STN excitation is incompatible with passive avoidance

In the preceding experiments, we tested the effects of STN excitation on CS signaled active avoids by applying optogenetic light during the active avoidance interval. To test the effect of STN excitation on passive avoidance, we used the AA3 procedure. During AA3, mice discriminate between two different tones (CS1 and CS2) delivered randomly; CS1 signals an active avoidance interval (like the CS in AA1/2), while CS2 signals a passive avoidance interval when mice are punished for crossing to the other compartment (ITCs are not punished). During AA3, mice must produce opposite actions to CS1 (active) and CS2 (passive). Thus, crossings (active avoids) during CS2 are errors.

In STN-ChR2 mice (n=11), we compared errors in control CS2 trials versus CS2+Light trials when STN excitation (40-66 Hz) is delivered during the passive avoidance interval. In later sessions, we also tested NoCS+Light trials, when the STN excitation occurs alone without consequence. Six of the mice evolved from CS1+Light experiments in Fig. 4, while the other five were naïve, and CS1+Light trials were not included in these experiments. The results did not differ and were combined. Mice discriminated between CS1 and CS2 trials by performing high rates of active avoids to CS1 and virtually nil to CS2, indicating their ability to passively avoid during CS2 (Fig. 5A; Tukey t(30)= 13.65 p<0.0001 CS1 vs CS2). However, STN excitation in CS2+Light trials blocked passive avoids causing mice to err (cross) on most of these trials (Fig. 5A; Tukey t(30)= 13.63 p<0.0001 CS2 vs CS2+Light). This occurred despite being punished for these errors. In fact, there was no difference in the rate of active avoids between CS2+Light trials and CS1 trials (85.2±2 vs 85.2±8; Tukey t(30)= 0.01 p=0.99 CS1 vs CS2+Light). Moreover, STN excitation alone in NoCS+Light trials drove the same high rate of active avoids as CS1 and CS2+Light trials (Fig. 5A; Tukey t(30)= 0.21 p=0.99 NoCS+Light vs CS2+Light), even though this did not predict the US and was not punished.

**Figure 5.**
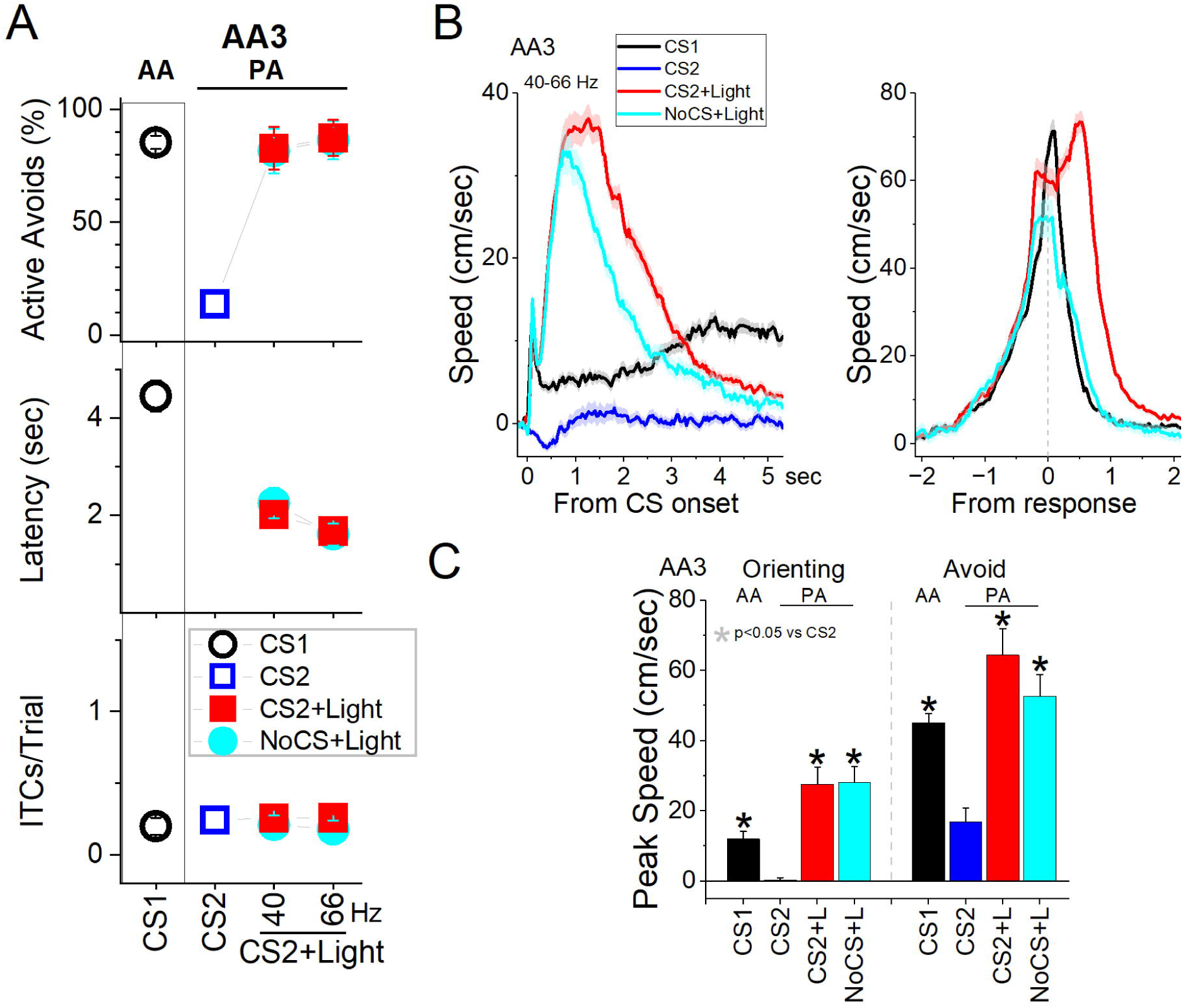
Optogenetic activation of ChR2-expressing STN glutamatergic neurons is incompatible with passive avoidance. ***A,*** STN excitation blocks signaled passive avoidance. In AA3, CS1 signals the animal to actively avoid, while CS2 signals the animal to passively avoid by not crossing. In addition, we tested CS2+Light trials and NoCS+Light trials (40-60 Hz light). STN excitation virtually abolished passive avoids to CS2 so that mice crossed despite being punished. Moreover, mice also crossed during NoCS trials when the light was delivered. ***B,*** Traces of overall speed during AA3 for the data in A. ***C***, Population measurements of peak speed for the data in B. Asterisks denote significant differences (p<0.05) between CS2 and the other conditions.

Movement tracking showed that the orienting response evoked by CS2, which can be small or even inhibitory (Zhou et al., 2023), was enhanced by STN excitation (Fig. 5B,C; Tukey t(30)= 11.62 p<0.0001 CS2 vs CS2+Light). The movement evoked by CS2 was also enhanced by STN excitation in association with the errors (Fig. 5B,C; Tukey t(30)= 11.79 p<0.0001 CS2 vs CS2+Light). In addition, because the mice are punished for crossing during CS2+Light trials, there was more movement during these trials related to the delivery of the punishment compared to NoCS+Light trials (Tukey t(30)= 3.92 p=0.04), which are not punished. This is noticeable by a hump in the speed trace of CS2+Light trials after the response error (red trace in Fig. 5C From response). Interestingly, when punished for these errors, mice move rapidly in reaction to the US but do not cross back to the other compartment, as indicated by the fact that the number of ITCs is virtually nil. These results indicate that STN excitation is incompatible with passive avoidance, and STN may serve as an important hub to regulate this inhibitory action.

In conclusion, STN optogenetic activation at 10-20 Hz drives normal signaled active avoidance responses, whereas STN activation >20 Hz shifts signaled active avoids earlier, and is incompatible with both cued inhibitory actions, such as passive avoidance, and the normal expression of caution when unsignaled actions are punished.

### STN excitation disables stopping

To determine whether mice can stop an already initiated action, we developed a Stop task in which mice perform two randomly interleaved trial types: regular active avoidance (AA) trials and Stop trials. Stop trials begin like AA trials with a CS_go_ (8 kHz), but when the mouse crosses an imaginary line positioned approximately one third of the distance toward the opposite compartment, a second cue (CS_stop_, 4 kHz) is presented instead, instructing the mouse to stop and refrain from crossing into the other compartment (Fig. 6A).

**Figure 6.**
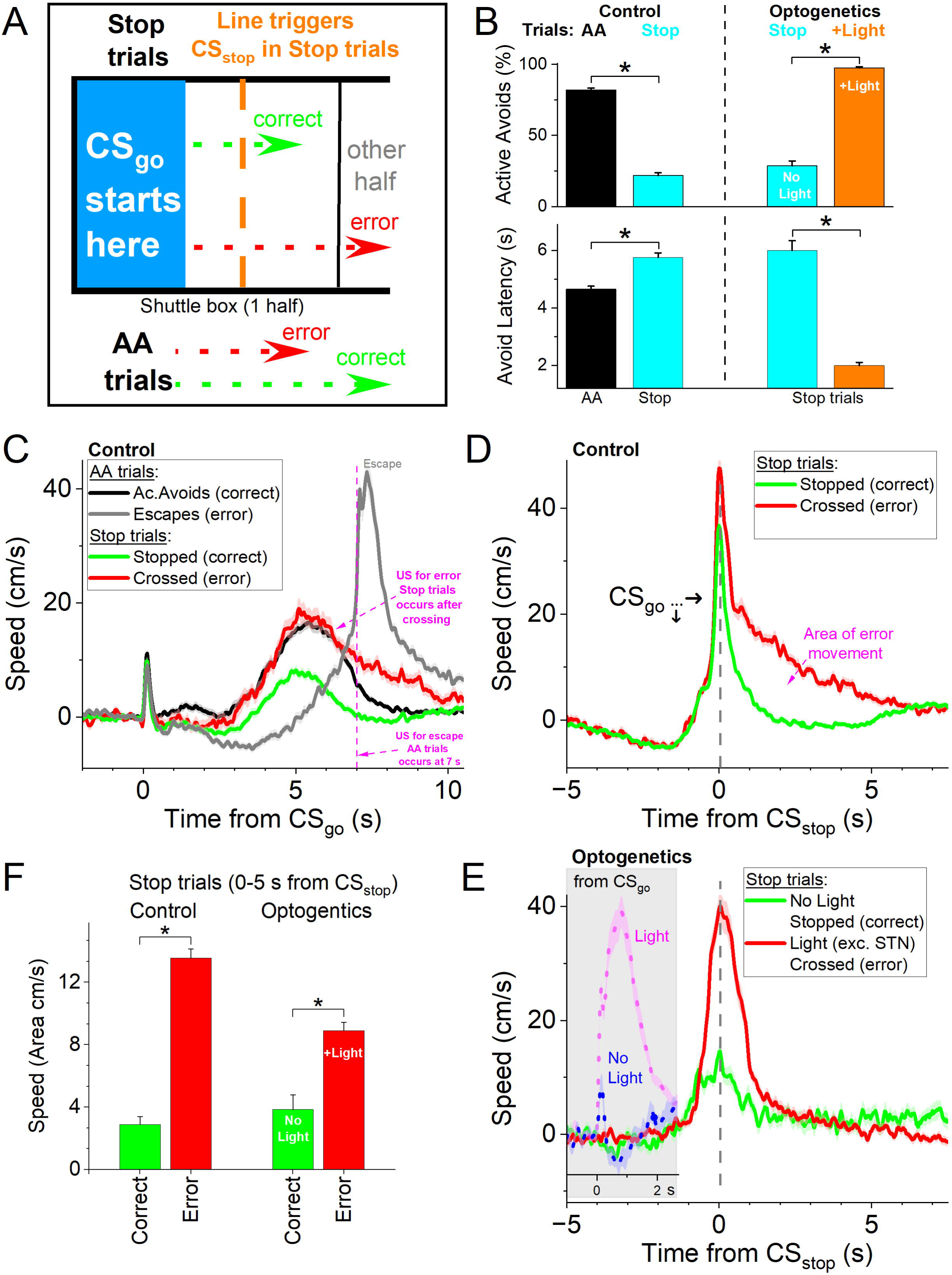
Optogenetic activation of STN neurons disrupts stopping. ***A,*** Schematic of the stop task consisting of regular active avoidance (AA) and stop trials. In AA trials, the regular CS_go_ is presented and mice must cross within the 7 s avoidance interval to avoid the US. Stop trials begin like AA trials with CS_go_, but when the mouse crosses an imaginary line toward the opposite compartment, a CS_stop_ is presented instructing the mouse to stop. If the mouse fails to stop and crosses into the other compartment, it receives a 0.5 s US punishment. ***B***, Percentage of active avoidance responses and avoidance latency in control mice comparing AA and Stop trials, and in optogenetic mice comparing Stop trials without and with STN excitation (40-66 Hz trains). The light delivery begins with CS_go_ at trial onset and continues through CS_stop_, until the mouse crosses or the trial ends. In Stop trials active avoids are errors. ***C***, Movement speed aligned from CS_go_ onset for AA and Stop trials in control sessions, separating correct trials and errors. ***D***, Control Stop trials aligned to CS_stop_, separated into correct trials and errors. Note the additional movement following CS_stop_ in error trials. ***E***, Optogenetic Stop trials aligned to CS_stop_, separated into correct Stop trials (No Light) and errors (Light). Inset (gray box) shows the same trials aligned to CS_go_. Note the rapid action initiation and failure to stop when STN excitation is delivered; mice fail to stop on nearly all light trials (97.3%). ***F***, Area under the movement curve for control and optogenetic Stop trials measured during the CS_stop_ interval (0-5 s).

Figure 6B (n = 11 mice) shows the percentage of active avoidance responses and avoidance latency during AA and Stop trials in control sessions (left). In Stop trials, active avoids are classified as errors. Mice reliably learned to stop their initiated action when instructed, resulting in a marked reduction in active avoidance responses during Stop trials compared to AA trials (t(240)=37.11 p <0.0001), and the latencies of the stop errors were longer than the those of correct active avoids (Fig. 6B lower; t(229)= 8.45 p < 0.0001).

In control sessions, speed traces aligned to CS_go_ onset during AA trials (Fig. 6C) reveal an initial orienting response followed by a delayed initiation of movement in correct active avoidance trials (black trace), whereas escape trials (gray trace) show a much longer delay before movement onset. During Stop trials in the same sessions, errors (red trace) exhibit a movement initiation pattern similar to active avoids, followed by additional movement associated with crossing and contingent punishment. In contrast, correct stops (green trace), show a shallower rise in speed reflecting curtailed movement after the stop instruction. Because the timing of CS_stop_ depends on when the mouse crosses the stop line, Stop trials are best evaluated when aligned to CS_stop_ onset. When aligned in this manner (Fig. 6D), correct Stop trials show a rapid suppression of movement following CS_stop_, whereas error trials show continued movement into the opposite compartment and punishment-evoked activity (correct vs error during Stop interval: t(190) = 19.45 p < 0.0001; Fig. 6F left).

We next tested the effect of optogenetic excitation of STN neurons during Stop trials. In optogenetic sessions, blue light stimulation (40–66 Hz) was delivered beginning at CS_go_ onset and continued through CS_stop_ until the trial ended or the mouse crossed. Under these conditions, STN excitation profoundly disrupted stopping: mice failed to stop on nearly all Stop trials (97.3%; % Avoids control vs light: t(113)=32.27 p <0.0001), instead rapidly initiating and completing the crossing, typically within 2 s from CS_go_ onset, despite presentation of CS_stop_ and contingent punishment (Fig. 6B,E;Avoid latency control vs light: t(102)=19.55 p <0.0001).

Movement aligned to CS_stop_ revealed a sharp increase in speed that was not curtailed by the stop signal, with speed profiles aligned from CS_stop_ resembling those aligned to CS_go_. Consistent with this, total movement during the CS_stop_ interval was significantly increased during STN excitation compared to control stop trials (Fig. 6F; t(105)=7.16 p < 0.0001).

Together, these results demonstrate that STN excitation not only disrupts passive avoidance but abolishes the ability to stop an already initiated action.

### STN excitation is not aversive

It has been proposed that STN stimulation might be aversive (Serra et al., 2023). Signaled active avoidance, in which a sensory cue predicts an aversive event, provides an ideal paradigm to test whether STN excitation is itself aversive and can function as a US. To address this, we trained naïve mice expressing ChR2 in STN neurons (with bilateral STN cannulas) using a modified AA1-3 procedure, where three CSs (CS1–CS3) were introduced from the first AA1 session. In this version, the US was STN excitation delivered as 66 Hz blue light (1-ms pulses) at ∼1.5 mW, titrated for each animal (n = 9) to reliably evoke rapid (<1.5 s) escape responses, comparable to those elicited by the standard aversive US. Control mice lacking opsin expression (No Opsin) were trained under the same conditions, including light delivery (n = 7).

During AA1, CS1 predicts the US light, while CS2 and CS3 predict nil. Crossing during CS1-3 turns off the CS and in CS1 avoids the light US. During AA2, intertrial crossings were punished by the light US (0.5 sec). During AA3, crossing during presentation of CS2 resulted in light US punishment (0.5 sec), while CS3 remained neutral. Despite the robust light-evoked escape responses during the US interval, ChR2-expressing mice failed to learn active avoidance to CS1 in AA1, AA2, or AA3 procedures (Fig. 7A). There was no difference in any of the measured parameters between the Opsin and the No Opsin groups (Mixed ANOVA Interaction GroupxCS; Tukey’s p>0.8 Opsin vs No Opsin during AA1-3). The CS1 cue, which predicted the STN excitation, did not increase avoidance behavior compared to CS2 or CS3, which were not predictive. In AA2, where intertrial crossings were also punished with STN excitation (0.5 sec), there was no reduction in these crossings. Similarly, during AA3, STN excitation failed to suppress crossings during CS2, which were also punished by the light. Mice had large numbers of crossings or spurious active avoids (∼45%) that were similar among the three CS’s despite the differences in the contingencies they predict, which is also indicative of normal exploratory behavior in an environment that is not punitive.

**Figure 7.**
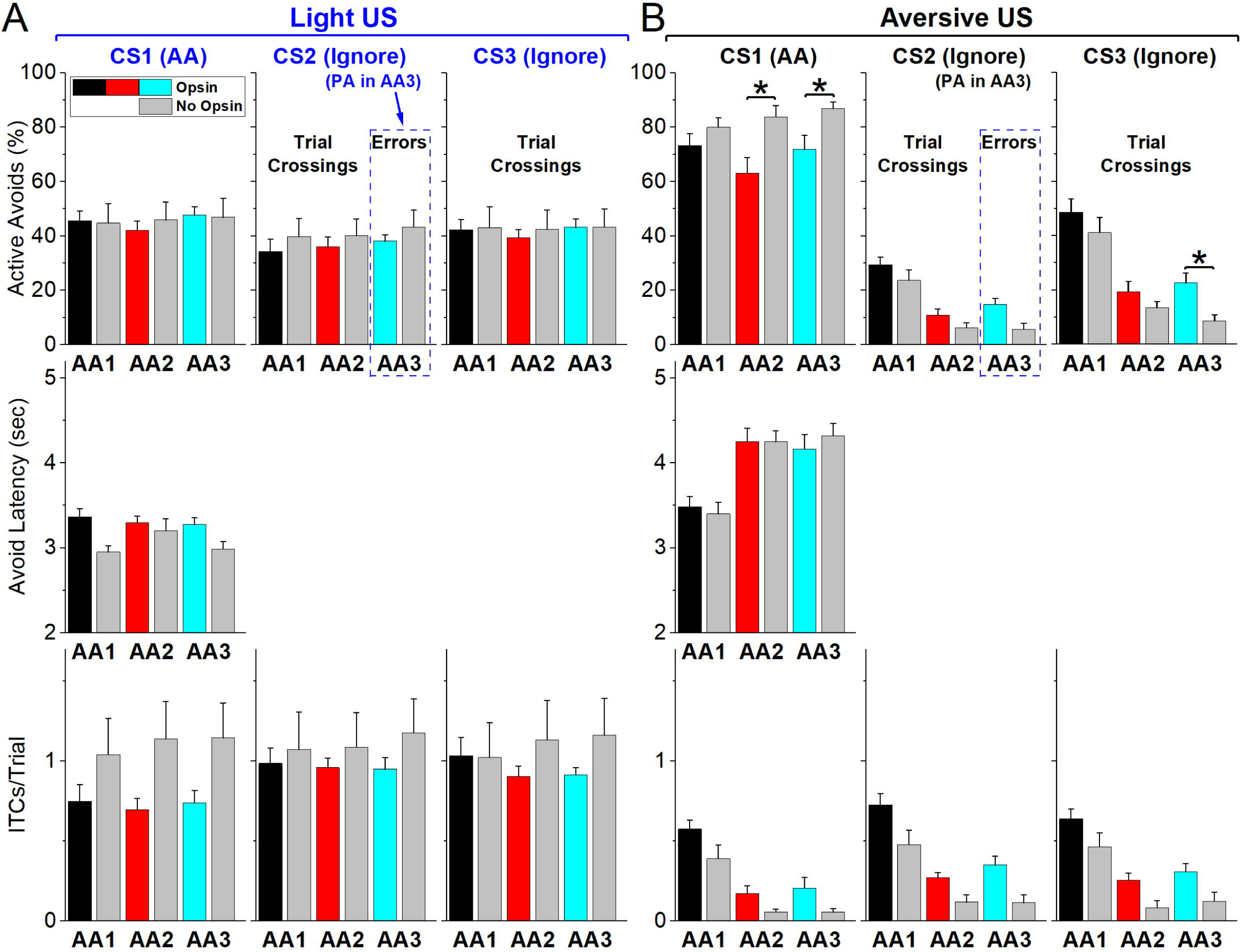
Optogenetic STN excitation is not aversive. ***A***, Signaled active avoidance procedures where the normal aversive US is substituted for STN optogenetic excitation that drives fast escape responses. Mice were subjected to AA1, followed by AA2 and then AA3 using 3 different CS’s. During AA1, CS1 predicts STN stimulation, while CS2 and CS3 predict nil. During AA2, ITCs are punished with STN stimulation. During AA3, mice must passively avoid by not responding when CS2 is presented, so active avoids driven by CS2 are errors punished with STN stimulation. The bars show side by side the results for Opsin (black, red, cyan) and No Opsin mice (gray) in the same procedures. Asterisks denote significant differences (p<0.05) between Opsin vs No Opsin mice. CS3 remains neutral through the AA1-2 procedures. ***B***, The mice in A are subjected to the same procedures using the normal aversive US. Asterisks denote significant differences (p<0.05) between Opsin vs No Opsin mice.

However, when the same mice were trained in the standard AA1 procedures using the regular aversive US (footshock and white noise), they rapidly learned to avoid during CS1 and significantly reduced crossings during CS2 and CS3—especially during AA2 and AA3, where these responses were punished (Fig. 7B). There was a small difference in the percentage of avoidance responses in Opsin vs No Opsin mice during AA2 and AA3 but not AA1 (Mixed ANOVA Interaction GroupxCS; Tukey q(28)=5.7 p=0.004 Opsin vs No Opsin during AA2; Tukey q(28)=4.3 p=0.044 during AA3) indicating that the mice had learned something about the CS1 predicting the STN excitation during the light US sessions, which reduced their avoids during the more cautious versions of the regular aversive tasks, perhaps emphasizing the role of STN in the more cautious responding that emerges during these sessions. In conclusion, these findings indicate that STN excitation, as applied in this study, does not function as an aversive US. Rather, the fast escape responses it elicits appear to result from its role in movement initiation, not from negative valence.

### STN excitation acts through midbrain projections

The results indicate that glutamatergic STN neurons have a critical role in signaled active avoidance. STN neurons have both ascending projections to the GPe area, and descending projections to SNr and the tegmentum in the midbrain (Kita and Kitai, 1987; Friedman and Yin, 2023; Prasad and Wallen-Mackenzie, 2024). Thus, we compared the effect of exciting the fibers of STN neurons ascending to GPe and descending in midbrain.

We prepared Vglut-STN-ChR2 mice but implanted the optical fibers in the GPe area (GPe group, n=7 mice), or in the SNr/tegmentum areas (Midbrain group, n=8 mice). The two sites in the midbrain were combined because the effects of exciting these areas were similar. A third combined No Opsin group (n=5 mice) had optical fibers implanted in the same targets as the other two groups. The locations of the optical fiber endings for these groups are depicted in Figure 8A.

**Figure 8.**
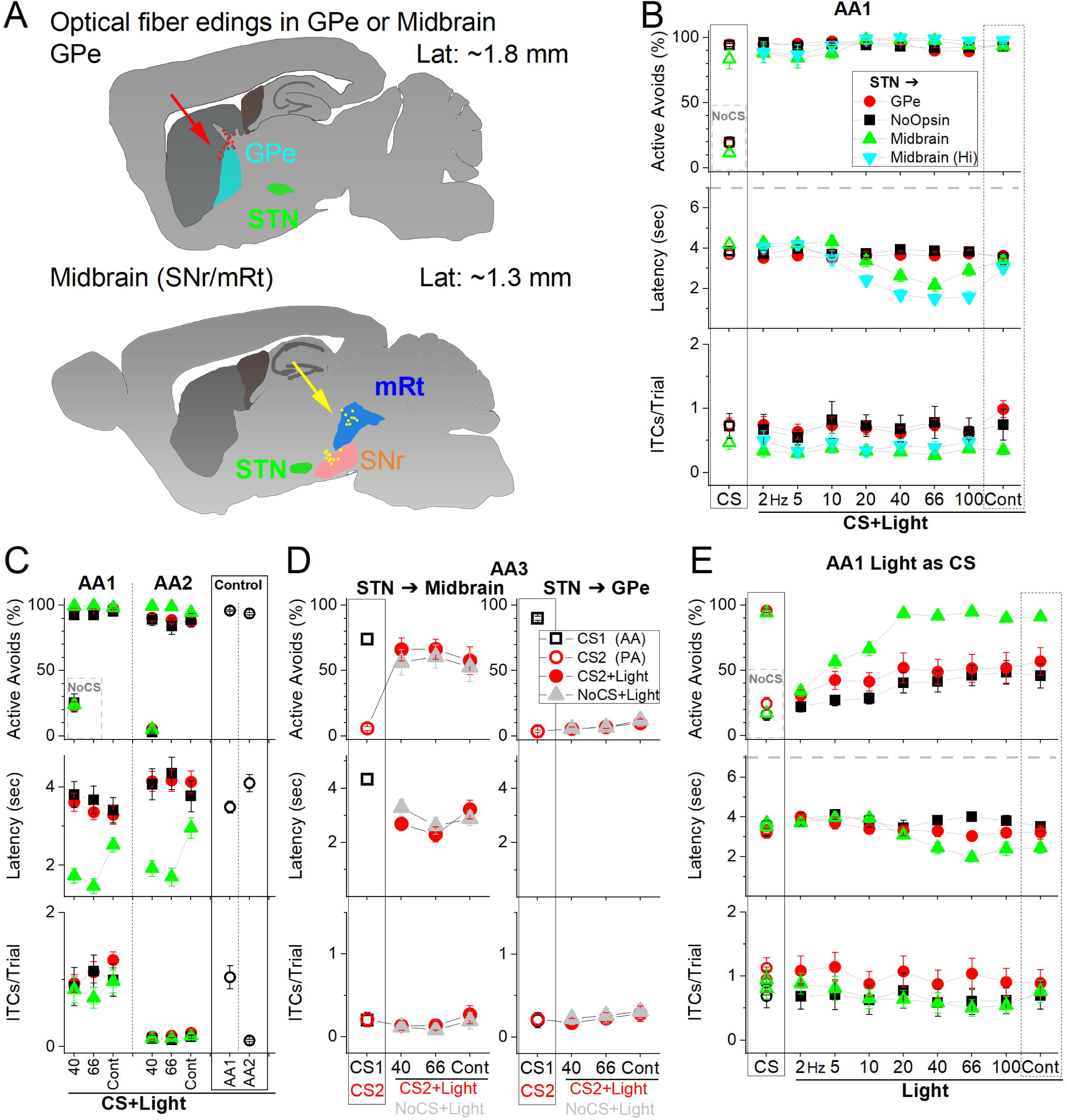
Optogenetic activation of STN glutamatergic pathways to the midbrain, not to GPe, mediate the effects of excitation within STN. ***A,*** Schematic of optical fiber locations for the midbrain targeting midbrain tegmentum (SNr and mRt combined) and GPe groups to target fibers originating in STN. ***B***, Effects of blue light patterns (CS+Light) applied in the Midbrain (filled triangles) or in the GPe (filled red circles) of STN-ChR2 mice on signaled active avoidance. A group of No Opsin mice are also included (filled back squares). Both optogenetic groups activated in the Midbrain and GPe received low and higher power light, but this is shown separately for the midbrain group as there was no effect of the light in the GPe group. Note that the effects in the Midbrain group occur for high frequency trains 20-100 Hz but not Cont light, which is consistent with the effective sustained activation of fibers with pulsed trains, not with Cont light. ***C***, Development of cautious responding from AA1 to AA2 (induced by punishing ITCs) is reflected in increased avoidance latencies and is suppressed by excitation of STN fibers in the midbrain, but not by excitation of STN fibers in the GPe or in No Opsin controls. Note the abolishment of ITCs during AA2 across groups, and the latency shift toward longer response times observed only in GPe and No Opsin mice. ***D,*** Signaled passive avoidance is blocked when STN fibers to midbrain are excited but not those to GPe or in No Opsin mice. In AA3, CS1 signals the animal to actively avoid (open black squares), while CS2 signals the animal to passively avoid by not crossing (open red circles). Application of CS2+Light (filled red circles) and NoCS+Light (filled gray triangles) trials (40-60 Hz light) in Midbrain but not in GPe virtually abolished passive avoids to CS2 and induced crossings in response to the light alone. ***E,*** Ability of STN fiber excitation in the Midbrain or GPe to serve as a CS assessed by comparing CS and Light trials. Only excitation of STN fibers in the midbrain was sufficient to drive active avoidance responses, whereas stimulation of STN fibers in the GPe was ineffective and comparable to responses in No Opsin controls.

We first explored if excitation of the STN pathways to GPe or midbrain drove earlier onset active avoid latencies characteristic of STN excitation. In CS+Light trials during AA1 (Fig. 8B; MixedAnova Group x Light), we found that excitation of STN fibers in the midbrain produced earlier onset latency active avoids than excitation of STN fibers in GPe (Tukey t(41)=5.2 p=0.001) or compared to the NoOpsin (Tukey t(41)=4.4 p=0.008) mice. The light patterns that were effective in driving earlier onset avoids were only the high frequency trains (40-100 Hz), which is consistent with these patterns being the most effective at activating fibers in a sustained fashion (Hormigo et al., 2019, 2021b, 2021a).

Next, we tested the development of caution reflected by the signaled action timing during the transition between AA1 and AA2 when all trials are CS+Light trials (40-66 Hz or Cont; Fig. 8C). We found that excitation of STN projections to the midbrain, but not those to GPe or in No Opsin mice, blocked the development of caution (Fig. 8C; MixedAnova Group x AA1/2). In all the groups, the ITCs were abolished during AA2. As occurs in control mice, in the No opsin (Tukey t(6)= 7.1 p=0.0024) and GPe (Tukey t(5)= 6.31 p=0.0066 AA1 vs AA2) groups the active avoid latencies shifted longer between AA1 and AA2. However, this shift did not occur in the midbrain group (Tukey t(7)= 1.5 p=0.3252) indicating that activation of the STN pathways to midbrain interferes with the development of caution when the unsignaled action is punished.

We then tested the effects of STN fiber excitation on signaled passive avoidance during AA3. Only excitation of STN pathways to the midbrain, but not those to GPe or the NoOpsin mice, blocked passive avoids to CS2 (Fig. 8D) by increasing the rate of errors to CS2 (Tukey t(63)= 7.81 p=0.0001 CS2 vs CS2+Light) to levels that were equivalent to CS1 (Tukey t(63)= 2 p=0.98 CS1 vs CS2+Light) despite the fact that these CS2 errors are punished. Moreover, the increase in crossings caused by activating STN pathways to midbrain also occurred in NoCS+Light trials, which were not different compared to CS2+Light trials (Tukey t(63)= 0.78 p=1 CS2+Light vs NoCS+Light).

Finally, we explored if these pathways could serve as an effective CS to drive active avoids in the absence of a natural auditory stimulus. In Light trials during AA1, we found that excitation of STN projections to the midbrain, but not those to GPe, were able to drive high rates of active avoids equivalent to the natural CS (Fig. 8E; MixedAnova Group x Light). Thus, activation of STN projections to the midbrain (Tukey t(38)>7 p=0.0007) at either 20-100 Hz or Cont drove higher rates of active avoids compared to GPe. Moreover, there were no differences in active avoid rates between the GPe and midbrain projection groups when they were activated at the lower 2-10 Hz optogenetic frequencies or when a natural CS was used.

Taken together, these results indicate that activation of STN pathways descending to the midbrain are responsible for the effects we observed when glutamatergic STN neurons were excited.

### STN projections to the midbrain exert an ipsiversive bias

We began this study by showing that excitation within STN drives movement with an ipsiversive bias. Next, we tested whether optogenetic excitation of STN projections to the GPe (n =7) or midbrain (SNr/mRt; n=8) would causally bias movements toward contraversive or ipsiversive directions. During open-field exploration (Fig. 9), we delivered unilateral blue-light stimulation to STN axon terminals using either continuous or 1-ms pulses at 10, 20, 40, or 66 Hz for 2 s. Light trains were delivered at random intervals within the same session. Identical procedures were performed in no-opsin control mice (n = 5), which did not express ChR2.

**Figure 9.**
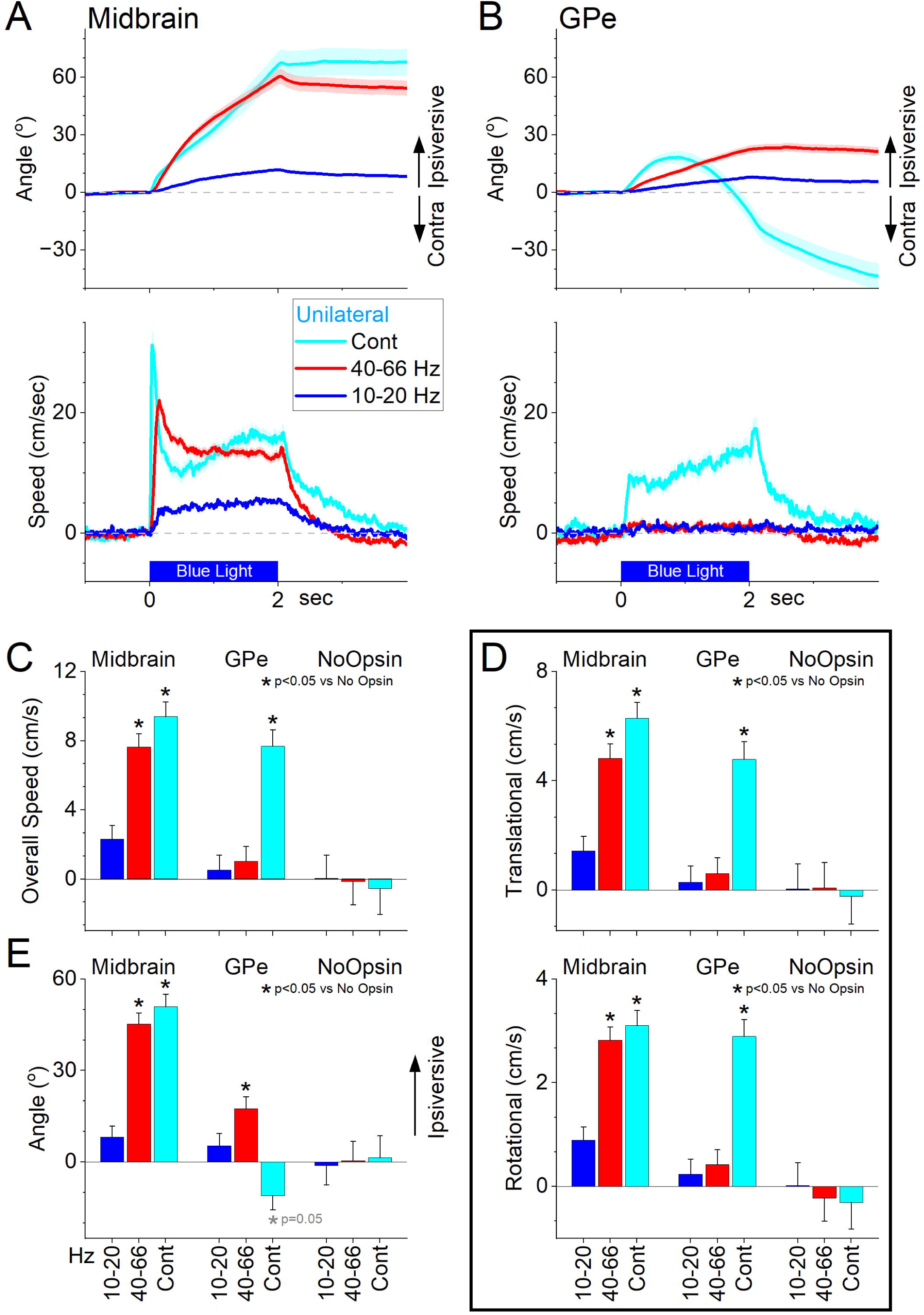
Optogenetic activation of the axons of ChR2-expressing STN neurons in the midbrain or GPe causes distinct movement biases. ***A***, Unilateral optogenetic excitation of STN projections in midbrain combining SNr and mRt locations drives an ipsiversive head orienting bias. Light patterns (2 sec) are delivered randomly as mice explore an arena. The top panel shows head orienting angle change from the onset of the light. The bottom panel shows the overall movement speed. Note the strong ipsiversive bias during 40-66 Hz and continuous excitation. ***B***, Same as A for GPe projections. ***C***, Population data comparing excitation of STN projections to midbrain or GPe with No Opsin mice on overall movement. Asterisks denote significant differences (p<0.05) between Opsin and No Opsin mice. ***D***, Same as C for translational and rotational movement components. ***E***, Same as C for peak angle.

Unilateral excitation of STN projections to the midbrain produced a robust ipsiversive head-orienting bias, with the magnitude of the effect scaling with stimulation frequency; continuous illumination produced effects comparable to high-frequency pulse trains (Fig. 9A). In contrast, excitation of STN projections to the GPe produced little evoked movement during trains, but continuous light evoked movement with a biphasic response, consisting of an initial ipsiversive turn followed by a pronounced contraversive turn (Fig. 9B).

Population analyses using mixed-effects models of overall movement (Fig. 9C), translational and rotational components (Fig. 9D), and movement angle (Fig. 9E) confirmed that stimulation of STN projections with high frequency light trains (40-66 Hz) to the midbrain reliably evoked movement with an ipsiversive bias compared to No Opsin controls (overall movement: t(47) = 5.0 p < 0.0001; angle peak: t(103) = 6.05, p < 0.0001), whereas trains of light applied to projections in GPe did not evoke more movement but biased ongoing movement ipsiversively (overall: t(48) = 0.7 p = 0.46; angle peak: t(58) = 2.3, p = 0.02). The ipsiversive bias caused by the high frequency trains was much stronger in midbrain compared to GPe (angle peak: t(59) = 5.0 p <0.0001). Continuous light revealed contrasting effects between STN targets, producing ipsiversive bias when stimulating midbrain projections (overall: t(70) = 5.8 p < 0.0001; angle peak: t(103) = 6.05, p < 0.0001) but a complex biphasic, contraversive dominated movement when stimulating GPe projections (overall: t(71) = 4.65 p < 0.0001; angle peak: t(107) = 1.95, p = 0.05). Finally, comparisons between animals with fibers placed in SNr versus mRt revealed no significant differences in movement measures.

Together, these results indicate that STN projections to the midbrain, including the SNr and mRt, promotes movement with a strong ipsiversive bias in a frequency-dependent manner. In contrast, STN projections to GPe produce contrasting effects that cause a contraversive bias when continuous light is applied.

## Discussion

Our findings demonstrate that STN excitation can function as an effective CS that elicits avoidance actions and precisely regulates action timing. Increasing stimulation frequency advances action onset in a graded manner, disrupting cautious responding without altering peak movement speed. Elevated STN activation also interferes with passive actions, rendering it incompatible with passive avoidance, or stopping initiated actions. Importantly, these effects are not driven by aversion but arise from stimulation-induced movement. We further show that these behavioral effects are mediated by STN projections to the midbrain, not to the GPe. Together with prior work establishing the necessity of STN–midbrain circuits for learned avoidance, these results show that STN activation can directly control the timing of cued, goal-directed action under threat.

### Ipsiversive lateralization

In our previous work, we found that ∼20% of STN neurons are strongly active during contraversive movements, whereas ∼25% exhibit a weaker ipsiversive bias (Zhou et al., 2025). In the present study, unilateral STN excitation in an open field reliably induced ipsiversive movements, with the strength of this bias scaling with stimulation frequency and intensity.

Notably, even studies that disagree on whether bilateral STN excitation facilitates movement (Friedman and Yin, 2023) or suppresses it (Guillaumin et al., 2021) consistently report that *unilateral* excitation produces ipsiversive turning.

Our results suggest that although STN excitation drives ipsiversive movements, the STN population encodes both directions, reflecting a distributed representation of ongoing and potential movement vectors. The ipsiversive bias during STN excitation only occurs when cells projecting to the midbrain (not those to GPe) are stimulated, which closely matches the effects of directly exciting its major downstream target, the GABAergic SNr, which also promotes ipsiversive turning under identical conditions (Hormigo et al., 2021a). A main pathway mediating this bias is the SNr to superior colliculus projection, since inhibiting the superior colliculus produces ipsiversive movements, whereas exciting it generates contraversive movements (Zhou et al., 2023). Thus, ipsiversive movements induced by STN excitation likely arise from STN-driven excitation of the SNr, which increases inhibition onto the superior colliculus and other orienting-related midbrain targets.

This framework supports the interpretation that STN activation engages a direct midbrain orienting circuit. However, while unilateral STN and SNr excitation produce similar ipsiversive directional biases, as discussed below, bilateral excitation of these nuclei yields distinct effects on cued actions, highlighting functional divergences between these areas.

### Effects of STN Excitation on Cued Actions

When examining the effects of STN excitation on behavior, our study includes two key features. First, we tested a wide range of optogenetic excitation patterns to modulate STN activity, uncovering differential effects that prior studies, often limited to a single stimulation pattern, may have missed. Second, we used a cued active avoidance task, where mice characteristically respond with slow onsets. This design enabled us to capture the impact of STN excitation at varying frequencies on action timing, and these effects could be overlooked in appetitive tasks due to the rapid onset of approach actions (Zhou et al., 2022).

Optogenetic STN excitation at medium frequencies (10–20 Hz) generated active avoidance responses closely resembling those triggered by a natural CS, matching in both timing and peak speed. In contrast, high-frequency excitation (>20 Hz) advanced action onset but did not increase peak speeds. Moreover, high-frequency stimulation evoked spontaneous crossings in untrained mice, suggesting an innate escape-like response to intense STN activation. In contrast, lower-frequency excitation did not induce spontaneous responses but effectively generated normal avoidance behaviors when STN activation served as a predictive CS for the US.

Evidence in humans shows that low-frequency oscillatory activity from the prefrontal cortex communicates with the STN to increase decision thresholds, promoting more deliberative, and thus slower, less impulsive, actions (Frank et al., 2007). In contrast, high-frequency STN DBS in Parkinson’s patients disrupts these signals, lowers decision thresholds, and induces impulsive decisions (Cavanagh et al., 2014; Herz et al., 2018, 2024; Pagnier et al., 2024). In agreement with these findings, we show that optogenetic stimulation of the STN in mice decreases goal-directed response latencies in a frequency-dependent manner. The correspondence of finding between mice and humans could enable STN excitation to be used as a model for studying impulsiveness, particularly in contexts when it leads to harmful consequences.

### STN excitation acts through the midbrain

Although both the SNr and midbrain tegmentum receive glutamatergic inputs from the STN (Kita and Kitai, 1987; Prasad and Wallen-Mackenzie, 2024), our loss- and gain-of-function results highlight an important asymmetry in how these downstream structures contribute to cued avoidance. In our previous work, direct excitation of SNr neurons potently suppressed active avoidance, whereas direct inhibition facilitated it (Hormigo et al., 2016). In contrast, here we show that excitation of STN neurons or their axons in SNr, which should excite SNr neurons, facilitate avoidance and trigger earlier action initiation. This functional mismatch indicates that the behavioral effects of STN excitation cannot be attributed solely to its influence on SNr output. Indeed, the facilitating effects on avoidance were also evoked by stimulating STN axons in the midbrain tegmentum in mRt. In a recent study, we also found that inhibiting STN neurons or its projections in either the SNr or the midbrain tegmentum abolished avoidance, while preserving escape responses elicited by the US (Zhou et al., 2025). Importantly, light delivered at the SNr is likely to affect STN fibers in route to the tegmentum, whereas light delivered directly to the tegmentum in mRt need not influence SNr-projecting fibers unless they are collaterals of those axons reaching mRt. Thus, the convergent effects of STN somatic and axonal modulation (inhibition and excitation) most parsimoniously implicate midbrain tegmentum targets as a critical locus through which the STN supports cued avoidance, which is consistent with the critical role of midbrain tegmentum in mediating cued avoidance (Hormigo et al., 2019).

Disambiguating the specific contributions of SNr-projecting versus tegmentum-projecting STN neurons will require future experiments designed to selectively isolate these pathways.

### STN excitation and aversiveness

Our findings align with STN’s purported role in action initiation in mice (Watson et al., 2021; Callahan et al., 2024), but the effects of high-frequency excitation may also reflect the aversive nature of excessive STN activation (Serra et al., 2023). Deciphering the exact sensation elicited by STN excitation in mice is challenging, particularly since even self-stimulation rewarding brain sites can become punishing or aversive under intense stimulation (Valenstein and Valenstein, 1964). A key distinction between escape behaviors driven by aversive stimuli and avoidance actions is the increased peak speed characteristic of escapes (Zhou et al., 2022, 2023; Hormigo et al., 2023). However, high-frequency excitation primarily shifted response onset earlier without increasing peak speed. This lack of increased peak speed suggests that the effects of STN excitation are not purely aversive.

To directly test whether STN excitation is aversive, we used high-frequency STN stimulation that induces fast escape responses –akin to an aversive US—as the US in signaled active avoidance procedures. If STN activation were aversive, it should substitute for the natural US and support conditioned avoidance. However, auditory cues that predicted STN excitation failed to drive avoidance responses, indicating that STN activation is not aversive. Notably, those same cues rapidly acquired the ability to drive avoidance when they were subsequently paired with a naturally aversive US.

Our results indicate that STN activation supports normal cued actions at lower levels of excitation (1-ms pulse trains <20 Hz), while higher excitation accelerates action onset in a manner that could be misinterpreted as aversive. These findings support a model in which STN activity promotes action initiation and fine-tunes the timing of goal-directed behaviors across contexts—from rapid-onset escape or approach responses to more cautious, slower-onset avoidance actions where risk evaluation, outcome prediction, and inhibition of premature responses are critical.

### Influence of STN during inhibitory and cautious behavior

An intriguing finding shows that high-frequency STN activation disrupts cued passive avoidance in the AA3 discrimination task, where CS1 signals Go (active avoidance) and CS2 signals NoGo (passive avoidance), resembling typical Go/NoGo procedures used in humans. When STN activation was applied during CS2, mice failed to inhibit their actions and responded to CS2 as though it were CS1. This suggests a direct mechanism by which STN activation can drive inappropriate responding in cued contexts, even when such responses lead to harmful outcomes. We also found that STN excitation interfered with the ability to stop actions that had already been initiated. Notably, similar disruptions of STN function may underlie pathological conditions in which inappropriate actions override goal-directed inhibition, as observed in mouse models of OCD (Parolari et al., 2021; Malgady et al., 2023). This may help explain the therapeutic efficacy of STN DBS in alleviating OCD symptoms in humans (Mallet et al., 2008).

An interesting aspect of our study is the impact of STN excitation on cautious behavior, specifically response timing when ITCs are punished (Zhou et al., 2022). The level of STN activation determined the timing of cued actions, with excessive excitation proving incompatible with the normal adjustment of response timing under punishment conditions. This suggests that heightened STN activity impairs behavioral flexibility and the ability to adapt in tasks requiring response inhibition. These findings are consistent with the strong encoding of cautious responding in the same behavioral tasks (Zhou et al., 2025), and also align with the proposed role of STN in action slowing and cautiousness under conditions of conflict or task difficulty in humans (Frank et al., 2007; Cavanagh et al., 2014; Herz et al., 2024). Our results indicate that when STN activity is elevated to high levels, subjects lose the ability to modulate action timing in response to behavioral demands. This has important implications for understanding action control in both typical and pathological states, underscoring the potential of STN as a critical target for regulating action timing in neurological and mental health disorders.

## Materials and Methods

### Experimental Design and Statistical Analysis

The methods used in the present paper were like those employed in our previous studies (Hormigo et al., 2023; Zhou et al., 2025). All procedures were reviewed and approved by the institutional animal care and use committee and conducted in adult (>8 weeks) male and female mice. Most experiments used a repeated-measures design in which each mouse or cell served as its own control (within-subject), but we also compared experimental groups between (between-group comparisons). To test the main effects of experimental variables, we used either repeated-measures ANOVA or linear mixed-effects models. Models included fixed effects (e.g., CS, light patterns) and their interactions. Random effects were specified as sessions nested within subjects, following the general syntax: variable ∼ (Factor1 * Factor2 * …) + (1 | Subject/Session) in lme4. Separate models were fit for each measure. Model fit was assessed by likelihood ratio tests comparing nested models, confirming that interactions improved explanatory power. For significant main effects or interactions, post-hoc pairwise comparisons were performed using *emmeans* with Tukey’s correction (Holm-adjusted).

To enable rigorous approaches, we maintain a centralized metadata system that logs all details about the experiments and is engaged for data analyses (Castro-Alamancos, 2022).

Moreover, during daily behavioral sessions, computers run experiments automatically using preset parameters logged for reference during analysis. Analyses are performed using scripts that automate all aspects of data analysis from access to logged metadata and data files to population statistics and graph generation.

### Strains and Adeno-Associated Viruses (AAVs)

To excite glutamatergic STN neurons using optogenetics, we injected AAV5-EF1a-DIO-hChR2(H134R)-eYFP (UPenn Vector Core or Addgene, titers: 1.8x10^13^ GC/ml by quantitative PCR) in the STN of Vglut2-cre mice (STN-ChR2 mice). No-Opsin controls were injected with AAV8-hSyn-EGFP (Addgene, titers: 4.3x10^12^ GC/ml by quantitative PCR) or nil in the STN. For optogenetics, we implanted dual optical fibers bilaterally in the STN or its projection targets. All the optogenetic methods used in the present study have been validated in previous studies using slice and/or in vivo electrophysiology (Hormigo et al., 2016, 2019, 2021b, 2021a).

### Surgeries

Optogenetics experiments involved injecting 0.2-0.4 µl AAVs per site during isoflurane anesthesia (∼1%). Animals received carprofen after surgery. The stereotaxic coordinates for injection in STN are (from bregma; lateral from the midline; ventral from the bregma-lambda plane in mm): 2.1 posterior; 1.7; 4.2. In these experiments, a dual (200 µm in diameter for optogenetics) optical fiber was implanted unilaterally or bilaterally during isoflurane anesthesia. The stereotaxic coordinates for the implanted optical fibers (in mm) are: STN (2-2.1 posterior; 1.5; 4.2-4.3), midbrain (3.3-3.7 posterior; 1.5; 2.9-4.1), and GPe (0.5 posterior; 2; 3.2). The coordinate ranges reflect different animals that were combined because the coordinate differences produced similar effects. No Opsin mice were implanted with cannulas in STN or its projections sites and the results were combined after confirming that light produced similar effects in these animals.

### Active Avoidance tasks

Mice were trained in a signaled active avoidance task, as previously described (Hormigo et al., 2016, 2019). During an active avoidance session, mice are placed in a standard shuttle box (16.1" x 6.5") that has two compartments separated by a partition with side walls forming a doorway that the animal must traverse to shuttle between compartments. A speaker is placed on one side, but the sound fills the whole box and there is no difference in behavioral performance (signal detection and response) between sides. A trial consists of a 7 sec avoidance interval followed by a 10 sec escape interval. During the avoidance interval, an auditory CS (8 kHz 85 dB) is presented for the duration of the interval or until the animal produces a conditioned response (avoidance response) by moving to the adjacent compartment, whichever occurs first. If the animal avoids, by moving to the next compartment, the CS ends, the escape interval is not presented, and the trial terminates. However, if the animal does not avoid, the escape interval ensues by presenting white noise and a mild scrambled electric foot-shock (0.3 mA) delivered through the grid floor of the occupied half of the shuttle box. This unconditioned stimulus (US) readily drives the animal to move to the adjacent compartment (escape response), at which point the US terminates, and the escape interval and the trial ends. Thus, an *avoidance response* will eliminate the imminent presentation of a harmful stimulus. An *escape response* is driven by presentation of the harmful stimulus to eliminate the harm it causes. Successful avoidance warrants the absence of harm. Each trial is followed by an intertrial interval (duration is randomly distributed; 25-45 sec range), during which the animal awaits the next trial. We employed four variations of the basic signaled active avoidance procedure termed AA1, AA2, and AA3.

In AA1, mice are free to cross between compartments during the intertrial interval; there is no consequence for intertrial crossings (ITCs).

In AA2, mice receive a 0.2 sec foot-shock (0.3 mA) and white noise for each ITC. Therefore, in AA2, mice must passively avoid during the intertrial interval by inhibiting their tendency to shuttle between trials, termed intertrial crossings (ITCs). Thus, during AA2, mice perform both signaled active avoidance during the signaled avoidance interval (like in AA1) and unsignaled passive avoidance during the unsignaled intertrial interval.

In AA3, mice are subjected to a CS discrimination procedure in which they must respond differently to a CS1 (8 kHz tone at 85 dB) and a CS2 (4 kHz tone at 75 dB) presented randomly (half of the trials are CS1). Mice perform the basic signaled active avoidance to CS1 (like in AA1 and AA2), but also perform signaled passive avoidance to CS2, and ITCs are not punished. In AA3, if mice shuttle during the CS2 avoidance interval (7 sec), they receive a 0.5 sec foot-shock (0.3 mA) with white noise and the trial ends. If animals do not shuttle during the CS2 avoidance interval, the CS2 trial terminates at the end of the avoidance interval (i.e., successful signaled passive avoidance).

When AA1–AA3 procedures were used to test the aversiveness of optogenetic stimulation, several modifications were introduced. Optogenetic light replaced the footshock and white noise as the unconditioned stimulus (US), and the tone continued during the US period.

Three conditioned stimuli (CS1–CS3) were included starting in AA1. CS1 and CS2 retained the same contingencies as in the standard AA procedures. CS2 and CS3 were neutral during AA1 and AA2, while CS3 remained neutral in AA3; neutral CSs were presented for 7 s and terminated if the animal crossed.

There are three main variables representing task performance. The percentage of active avoidance responses (% avoids) represents the trials in which the animal actively avoided the US in response to the CS. The response latency (latency) represents the time (sec) at which the animal enters the safe compartment after the CS onset; avoidance latency is the response latency only for successful active avoidance trials (excluding escape trials). The number of crossings during the intertrial interval (ITCs) represents random shuttling due to locomotor activity in the AA1 and AA3 procedures, or failures to passively avoid in the AA2 procedure. The sound pressure level (SPL) of the auditory CS’s were measured using a microphone (PCB Piezotronics 377C01) and amplifier (x100) connected to a custom LabVIEW application that samples the stimulus within the shuttle cage as the microphone rotates driven by an actuator controlled by the application.

### Stop task

The Stop task was used to test whether mice could interrupt an already initiated action in response to an instructive signal. Mice performed two randomly interleaved trial types: standard active avoidance trials and Stop trials. Stop trials began like regular active avoidance trials with presentation of a CS_go_ (8 kHz tone). If the mouse initiated an avoidance response and crossed an imaginary line positioned approximately one-third of the distance from the compartment door, a CS_stop_ (4 kHz tone) was presented instructing the mouse to stop and refrain from crossing into the other compartment for the next 5 s. If the mouse failed to stop and crossed into the other compartment during this interval, it received a 0.5 s US punishment. If the mouse successfully stopped, the CS_stop_ terminated after 5 s and the trial ended.

### Optogenetics

The implanted optical fibers were connected to patch cables using sleeves. A black aluminum cap covered the head implant and completely blocked any light exiting at the ferrule’s junction. Furthermore, the experiments occurred in a brightly lit cage that made it difficult to detect any light escaping the implant. The other end of the patch cable was connected to a dual light swivel (Doric lenses) that was coupled to a blue laser (450 nm; 80 mW) to activate ChR2. In experiments with ChR2 expression, blue light was delivered at constant power across different stimulation patterns, including continuous (Cont) and trains of 1-ms pulses at varying frequencies (2–100 Hz). Stimulation power was tested at several levels that were combined into two categories: low (0.5-2 mW) and high (3-6 mW). Power is regularly measured by flashing the connecting patch cords onto a light sensor—with the sleeve on the ferrule.

During optogenetic experiments that involve avoidance procedures, we compared different trial types: CS, CS+Light, Light, NoCS, NoCS+Light, and Light US trials. *CS trials* were standard avoidance trials specific to each procedure, without optogenetic stimulation.

*CS+Light trials* were identical to CS trials, except that optogenetic light was delivered simultaneously with the CS and the US during the avoid and escape intervals. *Light trials* were identical to CS+Light trials, but the CS was omitted to test whether optogenetic stimulation alone could substitute for the CS (cue). *NoCS trials* were catch trials with neither CS nor US, used to assess chance, baseline responses. *NoCS+Light trials* trials were identical to NoCS trials but included light stimulation to assess its effects in the absence of predictive cues or outcomes.

These differed from Light trials in that they did not signal or result in a US. *Light US trials* were like CS trials, but the US was optogenetic stimulation. To perform within group repeated measures (RM) comparisons, the different trial types for a procedure were delivered randomly within the same session. In addition, the trials were compared between different groups, including No Opsin mice that did not express opsins but were subjected to the same trials including light delivery.

### Video tracking

All mice in the study (open field or shuttle box) were continuously video tracked (30-100 FPS) in synchrony with the procedures and other measures. During open field experiments, mice are placed in a circular open field (10" diameter) that was illuminated from the bottom or in the standard shuttle box (16.1" x 6.5"). Optogenetic stimulation patterns were tested on movement and head direction bias during spontaneous exploratory behavior in sessions conducted on separate days from the behavioral task sessions, preceding and/or following completion of task testing. We automatically tracked head movements with two color markers attached to the head connector–one located over the nose and the other between the ears. The coordinates from these markers form a line (head midline) that serves to derive several instantaneous movement measures per frame (Zhou et al., 2023). Overall head movement was separated into *rotational* and *translational* components (unless otherwise indicated, overall head movement is presented for simplicity and brevity, but the different components were analyzed). Rotational movement was the angle formed by the head midline between succeeding video frames multiplied by the radius. Translational movement resulted from the sum of linear (forward vs backward) and sideways movements. *Linear* movement was the distance moved by the ears marker between succeeding frames multiplied by the cosine of the angle formed by the line between these succeeding ear points and the head midline. *Sideways* movement was calculated as linear movement, but the sine was used instead of the cosine. Pixel measures were converted to metric units using calibration and expressed as speed (cm/sec). We used the time series to extract window measurements around events (e.g., CS presentations). Measurements were obtained from single trial traces and/or from traces averaged over a session. In addition, we obtained the direction of the rotational movement with a *Head Angle* or *bias* measure, which was the accumulated change in angle of the head per frame (versus the previous frame) zeroed by the frame preceding the stimulus onset or event (this is equivalent to the rotational speed movement in degrees). The *time to peak* is when the *extrema* occurs versus event onset.

### Histology

Mice were deeply anesthetized with an overdose of isoflurane. Upon losing all responsiveness to a strong tail pinch, the animals were decapitated, and the brains were rapidly extracted and placed in fixative. The brains were sectioned (100 µm sections) in the coronal or sagittal planes. Some sections were stained using Neuro-trace. All sections were mounted on slides, cover slipped with DAPI mounting media, and all the sections were imaged using a slide scanner (Leica Thunder). We used an APP we developed with OriginLab (Brain Atlas Analyzer) to align the sections with the Allen Brain Atlas Common Coordinate Framework (CCF) v3 (Wang et al., 2020). This reveals the location of probes and fluorophores versus the delimited atlas areas.

## Acknowledgments

Supported by NIH grants to MAC.

